# Chronic recording of brain activity in awake toads

**DOI:** 10.1101/2024.10.16.618567

**Authors:** Daniel A. Shaykevich, Grace A. Woods, Lauren A. O’Connell, Guosong Hong

**Affiliations:** Department of Biology, Stanford University, Stanford, CA, USA; Department of Applied Physics, Stanford University, Stanford, CA, USA; Wu Tsai Institute for Neuroscience, Stanford University, Stanford CA, USA; Department of Material Science and Engineering, Stanford University, Stanford, CA, USA

**Keywords:** electrophysiology, mesh electrodes, *Rhinella marina*, cane toad, neural recording

## Abstract

**Background:** Amphibians represent an important evolutionary transition from aquatic to terrestrial environments and they display a large variety of complex behaviors despite a relatively simple brain. However, their brain activity is not as well characterized as that of many other vertebrates, partially due to physiological traits that have made electrophysiology recordings difficult to perform in awake and moving animals.

**New method:** We implanted flexible mesh electronics in the cane toad (*Rhinella marina*) and performed extracellular recordings in the telencephalon of anesthetized toads and partially restrained, awake toads over multiple days.

**Results:** We recorded brain activity over five consecutive days in awake toads and over a 15 week period in a toad that was anesthetized during recordings. We were able to perform spike sorting and identified single- and multi-unit activity in all toads.

**Comparison with existing methods:** To our knowledge, this is the first report of a modern method to perform electrophysiology in non-paralyzed toads over multiple days, though there are historical references to short term recordings in the past.

**Conclusions:** Implementing flexible mesh electronics in amphibian species will allow for advanced studies of the neural basis of amphibian behaviors.

## 1. INTRODUCTION

Since the discovery of “animal electricity” in amphibians more than 200 years ago (Piccolino, 1997), neural electrophysiology has greatly contributed to our modern understanding of brain function across animal life. Both intracellular and extracellular recordings, *in vivo* and *in vitro*, have allowed for biophysical characterization of neuronal activity, descriptions of brain responses to sensory stimuli, and examination of neural circuits underlying complex behavior (Cavanagh, 2019). Even as other methods for the live visualization of neuronal activity are popularized and new methods are developed (such as calcium imaging and opto-acoustic visualizations), much neuroscience research relies on traditional recording methodologies that may incorporate new technologies and genetic tools.

Advances of *in vivo* electrophysiology methods have led to recordings being implemented in increasingly difficult environmental situations, such as the wireless recording of brain recording in freely swimming fish (Vinepinsky et al., 2017) and very small Drosophilid flies (Swale et al., 2018). Yet, despite the initial discoveries leading to electrophysiology methodology taking place in frogs (Piccolino, 1997), limited modern neural electrophysiology work has characterized brain function in amphibians. This mostly includes measuring neural responses to sensory information, including visual processing in the retina or optic tectum (Buxbaum-Conradi and Ewert, 1995; Ewert et al., 1990; Finkenstädt and Ewert, 1983; Kühn and Gollisch, 2019) and auditory processing in the midbrain (Alluri et al., 2016; Hanson et al., 2016; Taylor et al., 2019; Wilczynski and Ryan, 2010). Historically, recordings from the amphibian telencephalon have been used to measure responses to sensory stimuli (Karamian et al., 1966; Kicliter and Ebbesson, 1976; Liege and Galand, 1972), but pallial brain regions are not commonly recorded from, likely due to low cell density and movement of the brain within the skull. This creates a large gap in our knowledge of amphibian brain function, as the forebrain is responsible for many aspects of cognition, including reward learning and spatial processing (Bingman and Muzio, 2017; Muzio et al., 1994; Papini et al., 2008; Sotelo et al., 2024). In addition, there has been no implementation of chronic recording in amphibians, and limited recordings in awake and moving animals, as amphibians are commonly paralyzed during recordings to reduce brain movement. What is needed to advance amphibian neuroscience research is the ability to (a) record from the amphibian telencephalon, (b) record from awake and moving animals, and (c) long term recordings that last more than a few hours.

In recent years, the development of new electrophysiological probes (including Neuropixels, NeuroGrid, and various electrode arrays) have improved the ability to record brain activity by upgrading spatial integration, temporal stability, and functional integration (Hong and Lieber, 2019). One class of novel devices are flexible mesh electronics (Dai et al., 2018; Fu et al., 2017, 2016; Hong et al., 2018; Lee et al., 2019; Woods et al., 2020), which are recording units designed to be more physically similar to neural tissue compared to rigid probes, allowing for longer recordings and reduced immune response. In rodents, flexible mesh electronics have allowed for chronic implantation that yielded recordings of local field potentials and single unit activity over the course of 8 months without probe repositioning (Fu et al., 2016).

We hypothesized that the tissue-like properties of flexible mesh electronics may help overcome issues of brain movement associated with recording neural activity in the amphibian telencephalon. Here, we demonstrate the use of a novel recording technology which allows us to record brain activity over multiple days in the telencephalon of the cane toad, *Rhinella marina* (Figure 1). Additionally, we demonstrate it is possible to record neural activity for up to 15 weeks. This methodology will allow for the more fine-scale study of behavior-related brain activity in amphibians, with the longer term goal of filling the taxonomic gap regarding amphibian brain function in vertebrate neural evolution.

**Figure 1.**
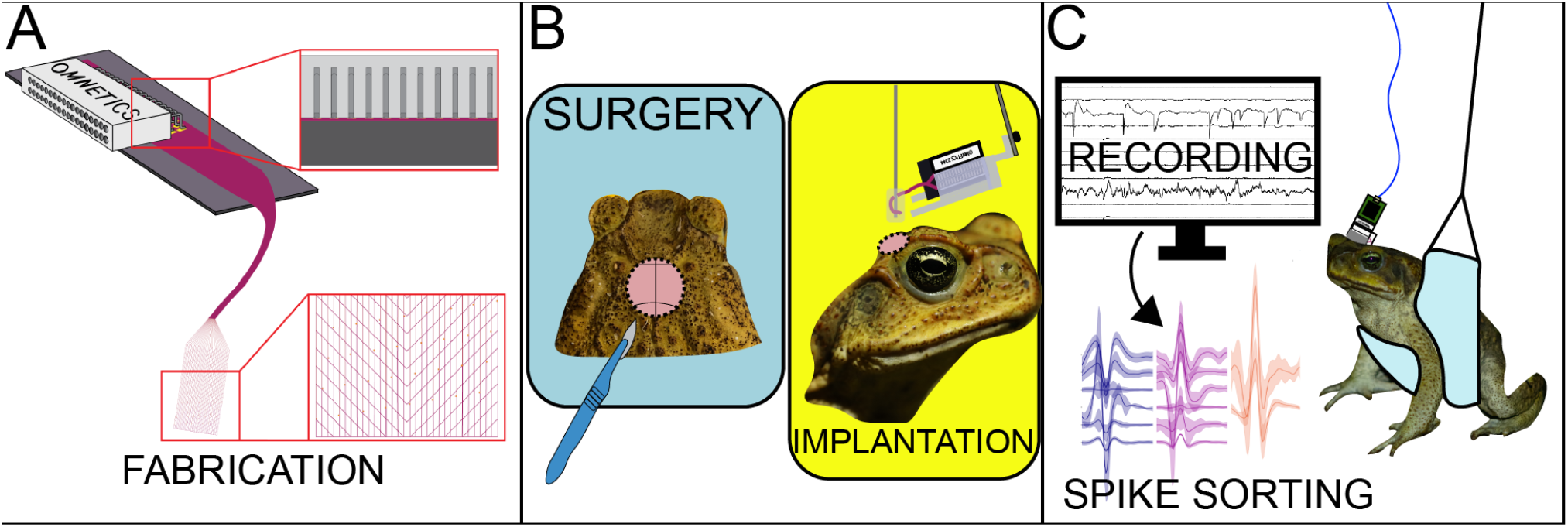
Project Overview. **(A)** Fabrication and preparation of flexible mesh electronic neural probes. **(B)** Cane toad surgery and implantation of mesh. **(C)** Chronic recordings in awake toads.

## 2. METHODS

### 2.1. Animals

All experimental subjects were sexually mature *Rhinella marina* collected in Oahu, Hawaii. Until surgery, toads were group-housed in glass terraria in a temperature and humidity controlled animal facility. Terraria were maintained at 20-25 °C and 80-100% humidity. Toads were fed gut-loaded crickets 3 times a week and had water dishes where they could rehydrate at any time.

### 2.2. Mesh Preparation

On-chip flexible mesh electronics were fabricated by the Hong Lab in Stanford University facilities as described in Woods (2023). Mesh devices were made up of 32 electrodes that had a lateral span of 2 mm. The fabricated device lies on a silicon substrate. Platinum electrodes are exposed on the end of passivated metal lines coated in SU-8, a photoresist polymer. All platinum electrodes are individually connected to Input/Output (I/O) pads at the other end of the mesh through gold interconnects. Each recording electrode is 20 μm in diameter (Woods, 2023).

Mesh were prepared as follows for implantation in *R. marina* (as depicted in Figure 2A). Fabricated mesh neural probes were bonded to Omnetics electronics connectors (A79024-001, Omnetics Connector Corporation, Minneapolis, MN, USA). First, photoresist was washed off from the device with acetone for ∼1 minute and then the device was rinsed in isopropyl alcohol. Next, using a stencil, Silver Conductive Epoxy Adhesive (MG Chemicals, Burlington, Ontario, Canada) was applied over the I/O pads and visually inspected under a stereoscope to check for shorts. Under stereoscopic guidance, an Omnetics connector was attached to a vacuum and lowered onto the I/O pads on a stereotaxic stage so that the connector legs aligned with the pads. Once the legs were contacting the pads, the epoxy was allowed to cure for roughly 20 minutes before the vacuum was lifted away, and the device was allowed to fully cure overnight.

**Figure 2.**
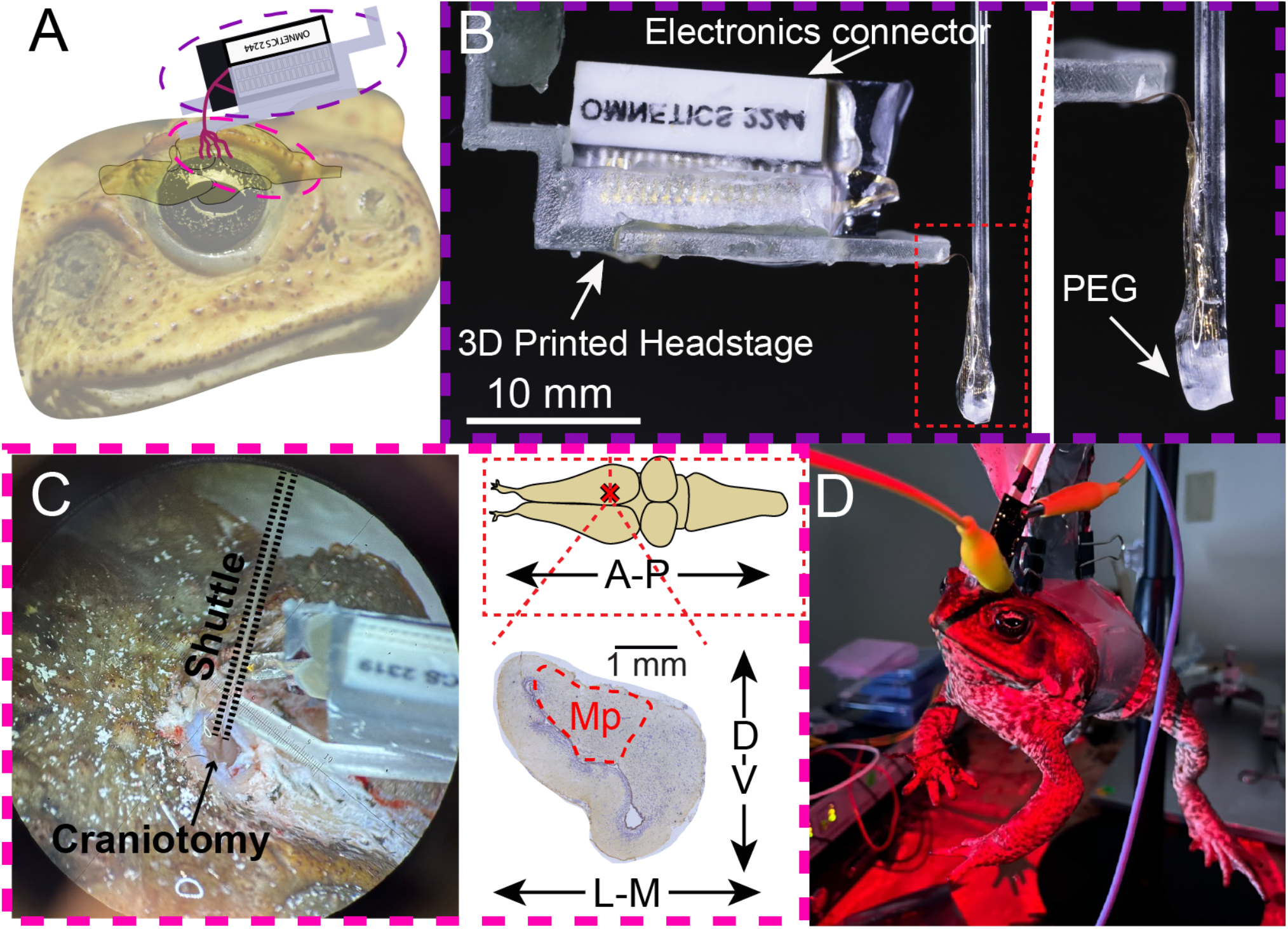
Implementation of flexible mesh electronics in cane toads. **(A)** Schematic of mesh implanted in toad brain. Colored ovals correspond to panels B and C. **(B)** Mesh bonded to the Omnetics connector and adhered into a 3D printed headstage. Inset shows mesh attached to glass shuttle with polyethylene glycol (PEG). **(C)** Implantation of mesh in a cane toad. Photograph taken through stereoscope of mesh on shuttle being lowered into toad telencephalon. Inset shows the position of implantation on the toad brain and a coronal section depicting the medial pallium (Mp). **(D)** Recording from an awake animal in a sling with implanted mesh connected to Intan RHX Data Acquisition system through the Omnetics connector.

The next day, dental cement was applied over the Omnetics legs and I/O pads, and over the lower edge of the Omnetics connector. When the surgery date approached, the mesh was etched from the substrate by suspending the device in etchant (Ferric Chloride 415, MG Chemicals, Ontario, Canada) by taping the substrate to the inner wall of a beaker and filling the beaker with etchant so that it covered the mesh, but did not reach the Omnetics Connector. Once the mesh had etched in ∼3-4 hours and could float off of the substrate, the device was removed from the etchant and excess was washed off with deionized (DI) H_2_O. The substrate was lightly scored with a diamond scribe below where the mesh was bonded to the Omnetics connector. The device was then fitted into a 3D printed headstage (seen in Figure 2B) and fixed with dental cement (C&B Metabond Quick Adhesive, Parkell, Edgewood, NY, USA). The mesh was floated off from the substrate in water and the substrate was snapped along the scoring lines. After the substrate was dry, dental cement was applied over the cleaved edge to prevent damage to the mesh. The headstage was taped to the wall of a beaker filled with DI H_2_O so the mesh floated and the Omnetics remained unsubmerged until the surgery. The day of or the day before the animal surgery, the headstage was screwed on to a shuttle adapter fitted to the arm of a stereotaxic stage. A glass capillary tube (Inner Diameter 0.15 mm, Outer Diameter 0.25 mm, Produstrial, Newton, NJ USA) was fitted into a pipette holder in the shuttle adapter and the tip of the shuttle was covered in polyethylene glycol (PEG 4000). Using a drop of water held on the end of a transfer pipette, the suspended mesh was attached to the shuttle via the PEG, and an additional drop of PEG was added to cover the mesh (Figure 2B).

### 2.3. Craniotomy and Implantation

At the beginning of the surgical procedure, the toad was placed into an MS-222 anesthetic bath (1.4 g/L) for about 20 minutes, or until the animal was unresponsive and did not respond to a toe pinch. The animal was removed from the bath and 4% Lidocaine was rubbed onto its skull and washed off with 1x phosphate buffered saline (PBS) after 3 minutes. A sterilized scalpel was used to remove all the skin from the snout to the back of the skull. Enamel etchant was applied to the exposed skull for 3 minutes and washed off with 1x PBS.

Prior to implanting the mesh, a craniotomy was performed in the middle of the skull in a line transecting the tympanum. A dental drill (Marathon-III, Saeyang Microtech, Daegu, Korea) with drill bits starting from 5/64” and transitioned to smaller sizes were used to drill through the skull. Surgical scissors and knives were used to cut through the dura over the brain. Once the brain was exposed, we visually confirmed that the telencephalon was visible. Once the craniotomy was complete, the toad was placed in the stereotaxic stage and the bonded mesh device, with the mesh attached to the shuttle, was loaded onto the stereotaxic arm. The shuttle was positioned over the medial pallium and lowered into the craniotomy opening until the end of the shuttle had penetrated 3 mm of brain tissue (Figure 2C). The shuttle was left in the brain for ∼10 minutes while the PEG dissolved. During this time, with the exception of toad M01, toads then had approximately 2 mm of 40 G stainless steel wire inserted into the telencephalon in the hemisphere opposite the mesh to serve as a physiological ground. After the mesh was fully released from the shuttle, the printed headstage was unscrewed from the shuttle adapter, the shuttle was retracted, and the headstage was attached to the skull with dental cement. Once the dental cement had set, any gaps and the exterior border were filled with Vetbond (3M, St. Paul, MN, USA). The section of the headstage that had attached the device to the shuttle adapter was broken off to prevent animals from using it to pry off the implant.

Following surgery, the toad was monitored until it had recovered from anesthesia and was fully mobile. It was housed alone in a glass terraria with automatic misting turned off and hydration provided through manual misting and a shallow glass dish with water. For 3 days post-surgery, the toad was subcutaneously injected with Meloxicam solution (OstiLox, VetOne, Boise, Idaho, USA) dosed at 0.4 mg/kg as an analgesic. Toads were monitored on a daily basis until recordings began. Following surgery, toads had a 2 week recovery period before any recordings were performed.

Implants occurred in 9 toads as described above. Three toads were recorded from over multiple days, as reported below. Two additional toads were recorded for 1 day and their implants fell out during the recording, after which the animals were euthanized. An additional 4 animals were implanted with mesh and their implants fell out prior to the 2 week recovery period being reached, after which the animals were euthanized.

### 2.4. Recordings

Toads were allowed to recover for 14 days post surgery before recordings began. Recordings were performed with the Intan RHD Recording System and Intan RHX Data Acquisition Software (Intan Technologies, Los Angeles, California, USA).

Recordings happened under two different regimes. Toad M01, a pilot toad with no physiological ground used to test mesh recordings prior to attempting them in awake toads, was recorded nine times over a period of 15 weeks, with recordings occurring once a week or once every two weeks. Prior to each recording, M01 was lightly anesthetized in a MS-222 bath (1.4 g/L). When anesthetized, a stainless steel wire was inserted into the edge of the toad’s beak to serve as physiological ground for the recording. This wire was connected to an alligator clip cable, the other end of which was clipped to a 2.54 mm mounting hole on the corner of the acquisition headstage. The Intan RHD 32 Channel Headstage, connected to the RHD Data Acquisition system by a 12-pin Omnetics cable, was plugged into the Omnetics connector of the implant. The recording happened as the animal was beginning to wake up, as evidenced by increased respiration and muscle tone, to increase the likelihood of capturing activity, as MS-222 is a sodium channel blocker that attenuates action potentials.

The remaining toads for which data is reported, F02 and F03, were recorded without anesthesia over the course of six recording sessions. Toads were put into a simple sling that held them around their midsection and were suspended from a desk-top stand. The Intan Headstage (wired to the RHD Data acquisition system) was plugged into the Omnetics connector of the implant. An alligator clip cable was used to connect the stainless steel physiological ground wire implanted in the toad to the 2.54 mm mounting hole on the corner of the headstage. Toads were recorded for ∼5 minutes every day. F02 was recorded for 5 consecutive days, and then a 6th day following a one day break. F03 was recorded for 6 consecutive days. However, as noise levels increased on the sixth days of recording, we did not consider the last days in our analysis.

### 2.5 Data Analysis

All raw recording data and filtered traces were plotted in Matlab for visual inspection to identify recordings with large movement artifacts or the presence of high noise levels. The recording files were then spike sorted using Kilosort4 (https://github.com/MouseLand/Kilosort) (Pachitariu et al., 2024) with manual curation in phy (https://github.com/cortex-lab/phy)(Rossant et al., 2016) to identify channels with single-and multi-unit activity. The bandpass frequency in Kilosort4 was set at 250 Hz. As Kilosort4 and phy are designed and optimized for dense electrode arrays with many contacts, we also used custom codes in our analysis. After identifying active channels and the amplitudes of the spikes, we used custom Matlab (v R2022b, MathWorks, Natick, MA, USA) code to visualize waveforms and interspike interval histograms to validate whether activity was likely to be single or multi-unit. A custom python script turned the sorted data from Kilosort4 and phy into a csv datasheet. R Studio (v 2023.09.1+494, Posit Software, PBC, Sunnyvale, CA) running R (v 4.3.1, R Foundation for Statistical Computing, Vienna, Austria) was used to create summaries of the results presented below.

### 2.6 Ethics Statement

All procedures were approved by the Institutional Animal Care and Use Committee of Stanford University (Protocol #33530). Toads were collected under the State of Hawaii, Department of Land and Natural Resources, Division of Forestry and Wildlife Protected Species (Permit No. WL21-16).

## 3. RESULTS

### 3.1 Recording from awake animals

We recorded two toads (F02 and F03) over five consecutive days while they were awake (not paralyzed and not anesthetized) (Figure 3). To assess our ability to measure single neuron activity over five consecutive days, we visually inspected raw recording data, performed spike sorting with Kilosort4 and phy to identify channels with single and multi-unit activity, and performed additional visualizations in Matlab.

**Figure 3.**
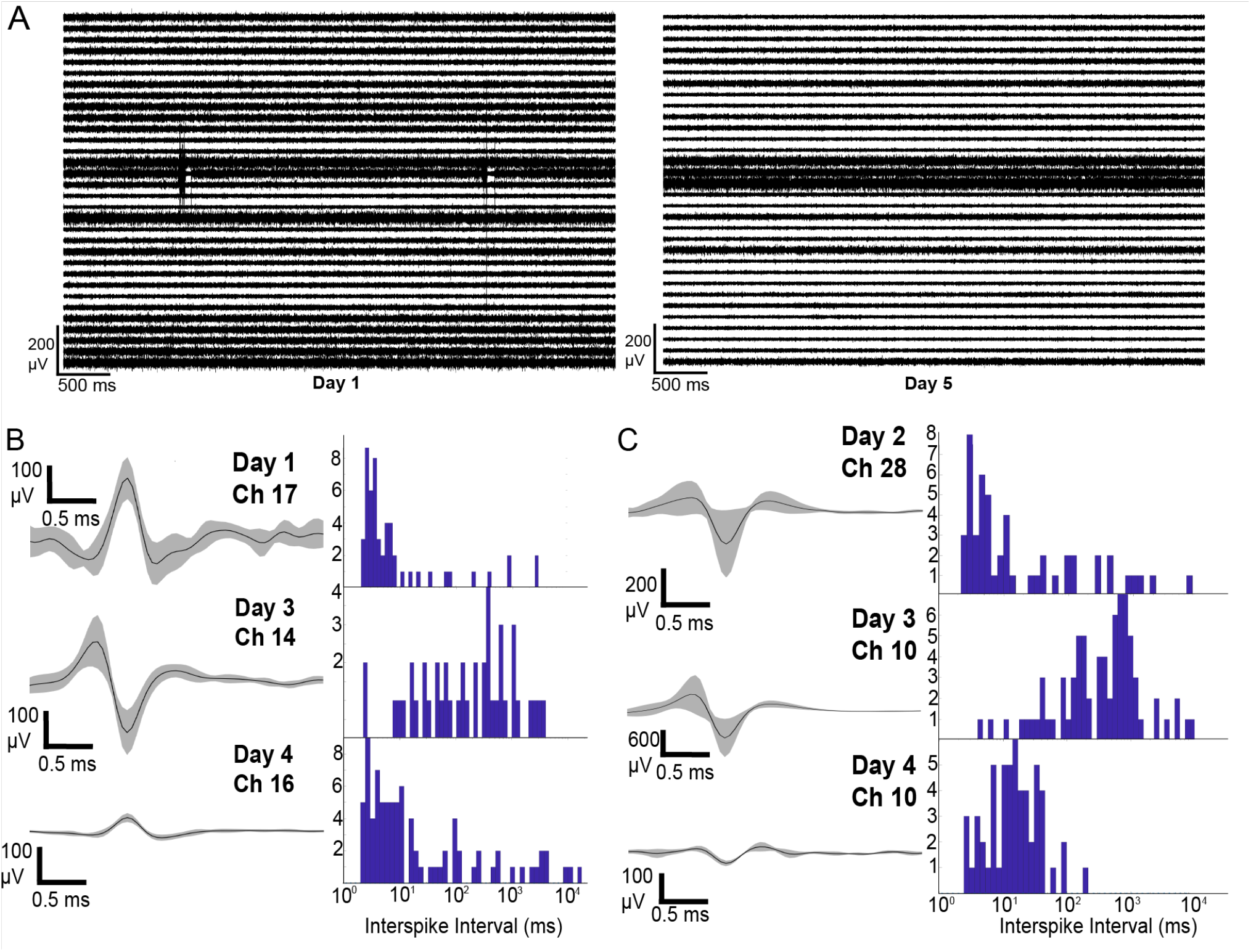
Chronic recordings in awake, partially restrained toads show single-unit activity. **(A)** Five seconds of high pass filtered (250-6000 Hz) traces from toad F02 taken on the first (left) and fifth (right) days of recordings. **(B-C)** Representative waveforms of single unit activity across recording days in (B) F02 and (C) F03, and interspike interval histograms for represented units. Lines represent mean waveforms and shading represents standard deviation of waveforms for single minutes of recording. Interspike interval histograms were generated from the same recordings.

For toad F02, we recorded 0-3 single units per day across all channels (mean 1.4) and 4-11 multi-unit signals per day for all channels (mean 6.4). The daily average amplitude of single units estimated by Kilosort4 ranged from 140-384 µV (231 ± 110 µV), which reflects the waveforms plotted by our Matlab spikesorter (Figure 3B). Amplitudes of multi-unit activity ranged from 187-533 µV (276 ± 146 µV).

For toad F03, we recorded 0-4 single units per day across all channels (mean 1.8) and 2-8 signals per day (mean 5.6) indicating multi-unit activity. The daily average amplitude of single units estimated by Kilosort4 ranged from 99-880 µV (472 ± 384 µV), also reflecting the waveforms plotted by our Matlab spikesorter (Figure 3C). Amplitudes of multi-unit activity ranged from 74-514 µV (236 ± 196 µV).

### 3.2 Recording from an anesthetized animal

We recorded from one toad (M01) over the course of about 15 weeks, where the animal was not freely moving but was anesthetized lightly with MS-222 during recording periods (Figure 4A). To assess our ability to measure single neuron activity across nine recordings, we inspected recording data visually, spike sorted in Kilosort4 and phy to identify active channels, and performed visualizations in Matlab.

**Figure 4.**
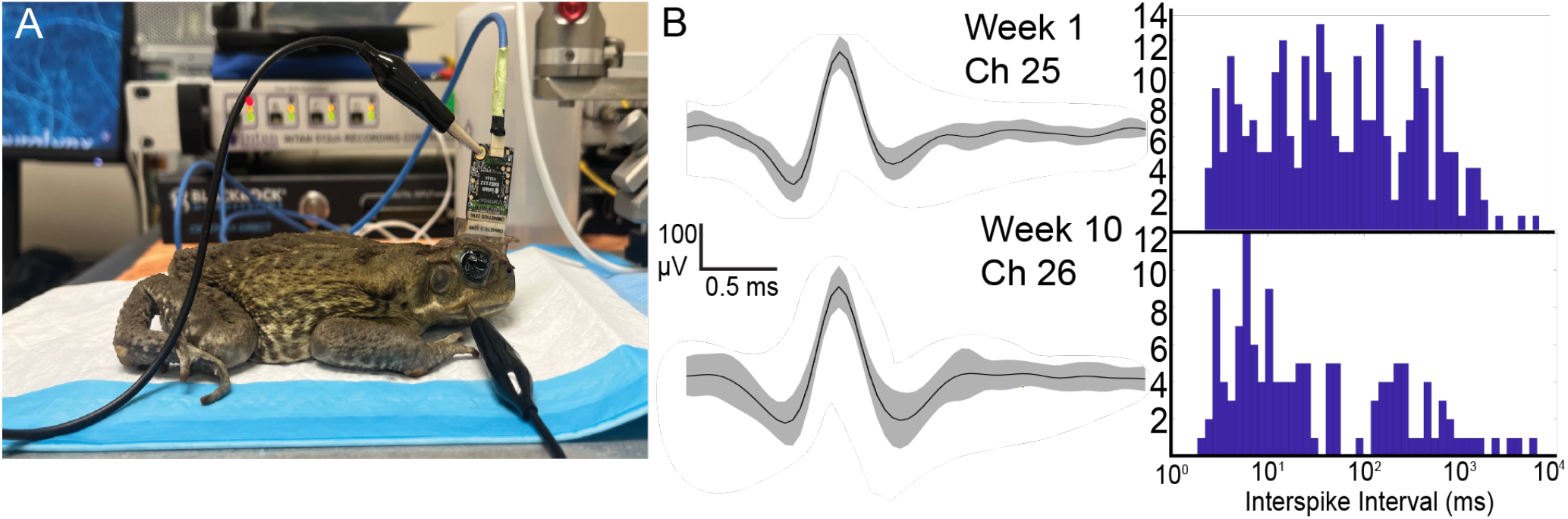
Mesh allows recording from an anesthetized toad over 15 weeks. **(A)** Toad M01 during recording, with Intan headstage connected to mesh and ground wire inserted in beak. **(B)** Representative waveforms of single unit activity in M01, with lines representing mean waveforms and gray shading representing standard deviation of waveforms, with corresponding interspike interval histograms.

In general, likely due to the less effective grounding in the animal, the activity recorded had a higher noise floor and was of much higher amplitude, and less single-unit and multi-unit activity was detectable in Kilosort4. However, on two days, we were able to record activity that appeared to be distinct, single unit activity (Figure 4B). In Kilosort4, the amplitude of these units was depicted at 1173 µV on the first day of recording and an amplitude of 1532 µV two months later. These amplitudes were likely disrupted by noise in Kilosort4, and appear at around 200 µV and 300 µV, respectively, in our Matlab visualization with thresholds applied. We recorded multi-unit activity on five of the nine recordings over 15 weeks, recording 1-3 signals in these days ranging in amplitude from 341-1345 µV (802 ± 368 µV).

### 3.3 Recording stability

To test recording stability over time, we compared activity on channels over recording days (Figure 5). We found the channel specific activity varied for all toads over the time spans shared, as active channels changed over the recording periods in all toads. In F02, activity (single-or multi-unit) was recorded on 21 channels over 5 days of recording: 10 channels showed activity on one day, 5 channels showed activity on 2 days, 5 channels showed activity on 3 days, and 1 channel showed recording on 4 days of recordings. In F03, 23 channels showed some activity over the course of the experimental period. Twelve channels showed activity on one day, 10 channels showed activity on 2 days, and 1 channel recorded some activity over 4 days of recording. In M01, we could distinguish activity on 8 channels. Six channels showed activity on one day, 1 channel showed activity on 2 days, and 1 channel showed activity on 3 days.

**Figure 5.**
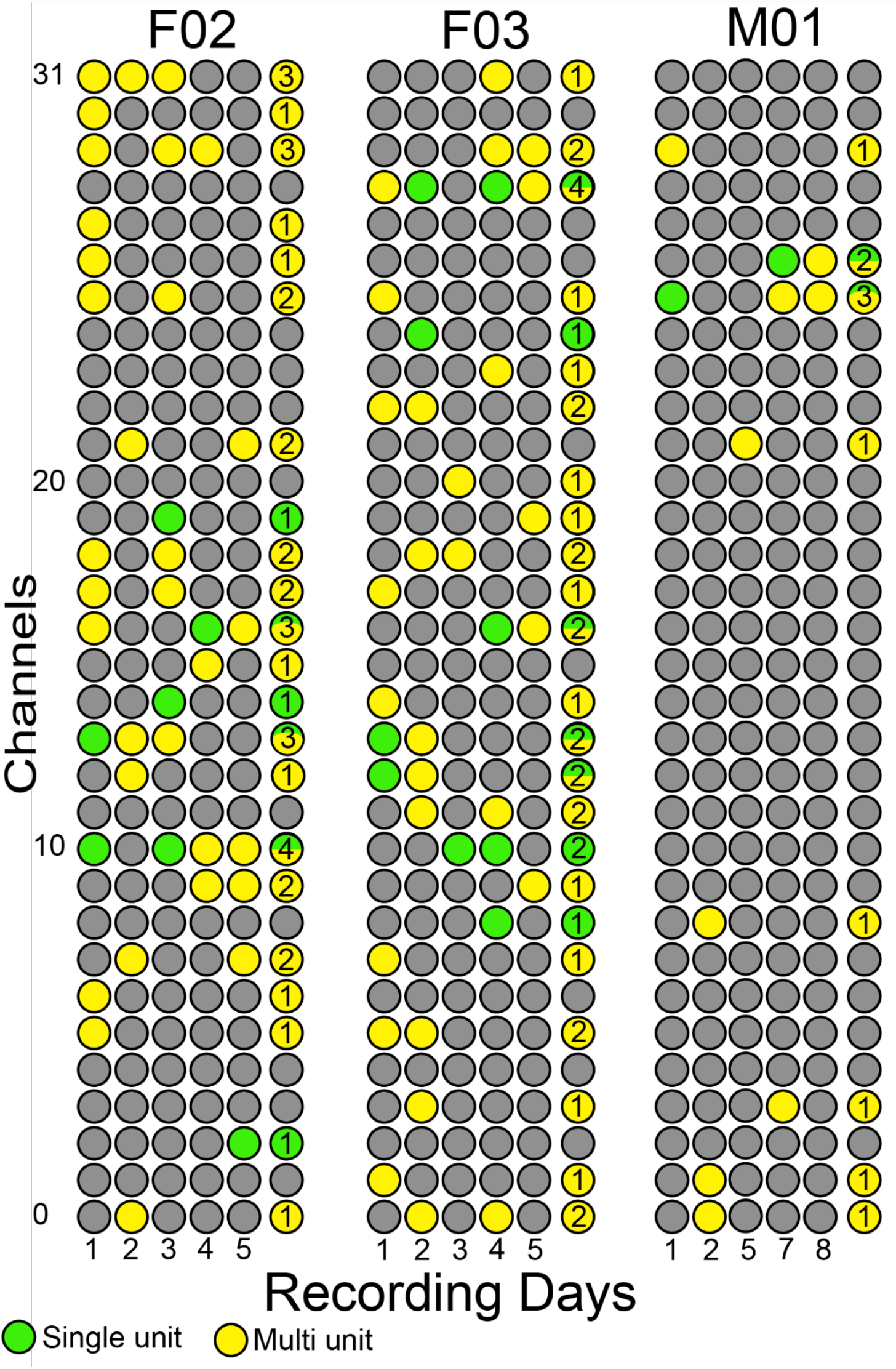
Stability of recording single- and multi-unit activity over recording days. Vertical axis depicts recording channels and the horizontal axis depicts days of recording for each of the three toads recorded over multiple days. Numbers in the last column for each toad indicate how many days a signal was recorded and the amount of green/yellow coloration depicts the proportion of signals that were single- or multi-unit. Five consecutive days of recording are depicted for F02 and F03; for M01, data shown represents the five recordings in which single- or multi-unit activity was detected, out of nine total recordings that occurred over fifteen weeks.

## 4. DISCUSSION

Here we demonstrate a method to record chronic extracellular activity from the amphibian telencephalon in awake, moving toads. This represents a major advancement in the experimental approaches available to study neural activity in amphibians by (a) allowing recording from a brain region that was previously incompatible with electrophysiology experiments, (b) allowing recording across several days, and (c) recording from animals that are not paralyzed. This presents a new opportunity for scientists to probe questions about neural function in amphibians. Further optimization is needed to increase yield of recordings and improve our ability to characterize action potentials. Overall, our results suggest that flexible mesh electronics are a useful tool to expand the possibilities of electrophysiology-based neuroscience research to new species.

### 4.1. Chronic electrophysiology in moving amphibians

The vast majority of neural recordings in amphibians occurs in paralyzed animals to overcome challenges in brain movement within the skull. Since at least the 1970’s, it has been commonplace to chemically immobilize amphibians with paralytics like pancuronium bromide and succinylcholine chloride (Ewert et al., 1990; Ewert and Hock, 1972; Leary et al., 2008; Womack et al., 2016). Indeed, chemical immobilization prevents the brain from moving, reducing noise during recordings and preventing damage to the brain. However, paralysis also reduces the possibility of what type of neuronal activity can be recorded during experiments, as the animal can only respond to stimuli presented to it and cannot perform behaviors. Thus, to study decision-making in amphibians, such as how they navigate their world or interact with other animals, recordings in freely moving amphibians is needed.

There have been many past attempts to perform electrophysiology in moving amphibians. There are records of extracellular recordings made from the optic tectum in freely moving toads using stainless steel electrodes decades ago (Laming et al., 1984; Laming and Ewert, 1983; Schürg-Pfeiffer, 1989). However, these *in vivo* recordings were regularly not stable, as Laming and Ewert (1983) report that recording from seven out of twelve freely moving toads could not be maintained for even 30 minutes in their experiment. More recently, a microdive allowed single recordings from awake Egyptian leopard toads (Mohammed et al., 2013), although we cannot locate additional studies utilizing this method. Overall, recordings from moving or unparalyzed amphibians are not reported in the last decade, where literature searches for “freely moving amphibian electrophysiology”, “mobile toad electrophysiology” and similar queries yield no modern results. Conversely, in other animals, including rodents, birds, and fish, it is commonplace to record activity from awake, moving animals to study neural activity related to mobile behaviors (Cohen et al., 2023; Payne et al., 2021; Yartsev and Ulanovsky, 2013). The ability to perform similar characterizations in amphibians will allow for a better understanding of the evolution of neural function across the vertebrate lineage.

Electrophysiological recordings in paralyzed animals generally do not allow for recordings lasting beyond 2 days, as the animals must be re-immobilized and the recording units lowered back into the brain. Implanted units in other vertebrates have already allowed for recordings that take place over days, weeks, and months, with flexible mesh electronics allowing for recordings happening over 8 months (Dai et al., 2018). Chronic recordings allow for the tracking of changes that happen over time and in response to experimental treatments. The ability to perform recordings over multiple days in amphibians may also benefit questions commonly asked in amphibians, such as responses to visual and auditory stimuli. Thus, implementing flexible mesh electronics in these studies may result in novel findings through the longitudinal observation of neural activity.

### 4.2. Recording telencephalic activity in amphibians

Most studies of electrophysiology in amphibians have concentrated on relatively few brain regions to characterize responses to sensory stimuli. For example, *in vivo* extracellular and whole cell recordings of the midbrain torus semicircularis, the anuran homolog to the mammalian inferior colliculus, have examined the duration and interval selectivity of neurons responding to sound stimuli (Alluri et al., 2016; Hanson et al., 2016). *In vivo* extracellular recordings from the optic tectum helped characterize how amphibians respond to objects moving in “worm” and “antiworm” orientation (Ewert, 1987; Schürg-Pfeiffer, 1989). However, there is a lack of modern *in vivo* electrophysiology methods for the amphibian telencephalon, even though this large segment of the brain holds regions involved in complex behaviors like reward learning (Muzio et al., 1994, 1993), parental care (Fischer et al., 2019), and spatial navigation (Shaykevich et al., 2024; Sotelo et al., 2022, 2019, 2016).

In the past, there was more research done about the function of the amphibian telencephalon. Kicliter and Ebbeson write that, prior to the 1970’s, research in this area either focused on the effects of ablations and stimulation on motor function, or studies using recordings or ablations to examine the role of the telencephalon in sensory discrimination (Kicliter and Ebbesson, 1976). These early recording studies occurred in paralyzed frogs and examined evoked potentials in response to visual and auditory stimuli (Karamian et al., 1966; Liege and Galand, 1972), similarly to the focus on sensory representation in modern amphibian electrophysiology. Outside of *in vivo* recordings, *in vitro* recordings from toad brain preparations have been used to describe the architecture of sensory input in a toad telencephalon (Laberge and Roth, 2007).

Past *in vivo* recordings in paralyzed frogs or recordings in slice preparations cannot characterize neural activity directly related to behavior, which is a major challenge in modern behavioral neuroscience in amphibians. However, there are a few reports of recordings in freely moving toads from the 1980’s that showed the promise of behaviorally relevant results with respect to responses to visual stimuli (Laming and Ewert, 1983; Schürg-Pfeiffer, 1989). The recordings from the telencephalon of awake toads demonstrated here open up new possibilities for research in this area. Furthermore, we were successful in recording for more than one day, which would further expand electrophysiology in amphibians by allowing for the study of changes in brain activity over the span of a few days, or even a few months. Our next steps will be to examine behaviorally-relevant neural coding with this methodology.

### 4.3 Further optimization of recording protocol

The recording protocol reported here needs further optimization before it can be implemented more widely. For example, we report activity recorded from one frog over the course of three months to show that it is possible for the implant to remain in a toad for that period of time. However, likely due to the lack of implanted physiological ground in this surgery (the toad was grounded through the mouth during recordings), the recordings were noisy and it was difficult to distinguish single units. In subsequent surgeries, with successfully implanted ground wires, we overcame this issue. After three months, the dental cement holding toad M01’s headstage came off as significant amounts of skin had regenerated under the cement. However, the fact the implant could say on for that period of time is extremely promising. Implants fell out prematurely in other toads as well, which may be partially attributable to the humid and wet conditions required for amphibian housing. We found that using enamel etchant improved the longevity of our implants, but additional measures, such as using bone screws, may be necessary to ensure these implants have a higher rate of success in the future.

The quality of the recordings and channel-specific activity varied from day to day (Figure 5). This suggests that, even with the mesh remaining implanted and activity detected from day to day, movement of the brain and shifting of the contacts in between recording periods may significantly alter the signal recorded across channels. Further experiments testing neuronal responses to stimuli and neuronal firing related to behavior are needed to test the feasibility of the chronic recordings in studying various types of neuronal activity.

We successfully managed to record from two toads over the course of 5 days with lower noise levels and a better ability to sort out activity (we stopped these experiments after 6 days of recordings, and the implants did not fall out in these two toads). Given that the device has 32 channels, we only ever saw partial yield of total recording capacity on any given day, seeing a maximum of 13 units on a day of recording. Low yield could be due to the low density of cells in the medial pallium and contacts not being near enough to neurons to pick up activity. Low yield could also be a result of poor bonding of the mesh with silver epoxy to the Omnetics connector, and better control of this manual step could lead to higher yield. Similarly, redesigning some aspects of the mesh fabrication to test more compact or more spread out distribution of contacts may result in a mesh better suited for recording from the toad telencephalon.

### 4.4 Expanding use of flexible mesh electronics

Flexible mesh electronics are an exciting technological development that has been implemented in neuroscience studies using rodent electrophysiology (Dai et al., 2018; Fu et al., 2016; Hong et al., 2018; Lee et al., 2019). Our results expand the utility of mesh electronics to a non-mammalian research organism. Our study shows that these recording units can not only be used in other species, but can potentially solve problems that prevented recordings from being achieved in the first place, such as sparse cells and large brain movements. Increasing the implementation of mesh in more model species will allow for greater optimization and further refinement of electrode architecture for different recording scenarios.

## 5. CONCLUSION

We demonstrated an extracellular recording protocol that allows for chronic recordings in awake toads with the use of flexible mesh electronics. This advancement will allow for a greater breadth of neuroscience in amphibians and provide methodology that can be used to study the neural activity underlying behaviors and responses to environmental stimuli. We hope to see this protocol adopted by other groups so that we can continue optimizations and broaden the understanding of neuroethology in frogs and other amphibians.

## Supporting information

Matlab scripts

## 6. ACKNOWLEDGEMENTS

We acknowledge that the laboratory portion of this research was conducted on the ancestral lands of the Muwekma Ohlone people at Stanford. We understand the implications of the historical and present colonialism the Ohlone people experience and celebrate their continued stewardship of their lands.

We would like to thank Julie Elie and Lisa Giocomo for consulting on the data analysis portion of this project. We would also like to thank Han Cui and Jack Daly-Bartley from the Hong Lab for additional support during various phases of the project.

## 7. DATA AVAILABILITY

Matlab scripts for spike sorting and visualizations have been uploaded as supplementary materials. Authors can provide raw recording files upon request.

## 8. AUTHOR CONTRIBUTIONS

Conceptualization: D.A.S., L.A.O, G.H.; Methodology: D.A.S., G.A.W., G.H.; Formal analysis: D.A.S., G.A.W.; Investigation: D.A.S., G.A.W.; Resources: L.A.O., G.H.; Data curation: D.A.S., G.A.W.; Writing – original draft: D.A.S.; Writing – reviewing and editing: L.A.O., G.H., G.A.W.; Visualization: D.A.S., G.A.W. ; Supervision: L.A.O., G.H.; Project administration: L.A.O., G.H.; Funding acquisition: D.A.S., L.A.O., G.H.

