## Supplementary material for "Chronic recording of brain activity in awake toads": Matlab scripts: Code_readme.rtf

This file contains the Matlab scripts that were used to analyze and visualize the data presented in “Chronic recording of brain activity in awake toads.” The following are brief descriptions of the included scripts/ 

spike_sorter_core_submission.m - This script allows you to set the parameters of spike sorting. It references read_Intan_512RHD_no_prompt.m, which can be dowloaded from the Intan website. 
The following parameters can be manipulated:
- startSortingTime and endSortingTime: allows you to pick a time range (in seconds) in a recording file to consider for spike sorting
- channels- allows you to select which channels to sort
- Time and amplitude constraints:
	-pre/post spike durations
	-detectMode: allows you to select “positive” peaks, “negative” peaks, or “both”
	-thresholdSetValue: allows you to set an amplitude value (in microvolts) that spikes must exceed to be sorted
	-artifactCutoff: Maximum amplitude for artifact removal in microvolts
-Bandpass filter parameters
	-samplerate - 20000 Hz for all recordings in this experiment
	-passBand - allows you to select frequency band; 250-6000 Hz for this experiment

spike_sorter_submission.m - This is the script you run to perform spike sorting, and it references the spike_sorter_core_submission.m script.  You can change the plotting parameters in this script. Daniel Shaykevich adjusted this script with the help of chatGPT, to plot mean waveforms with standard deviation shading, and checked the work to confirm it was accurate. Currently outputs filtered traces of sorted channels with peak indicators marking spike locations, a plot with all individual sorted waveforms overlayed on each other,  a plot of mean waveform of sorted spikes with standard deviation shading, and the raw traces from all filtered channels. 

ISI_log.m - This script plots the inter-spike interval histograms of sorted channels using the Spike_Locations.dat file output by running spike_sorter_submission.m. Daniel Shaykevich adjusted this script with the help of chatGPT to plot ISIs on a logarithmic time scale. 
