## Supplementary material for "Chronic recording of brain activity in awake toads": Matlab scripts: ISI_log.m

```
% This .m file takes the spike location files as the input and plots their ISI's.

% Written by Guosong Hong on July 7, 2015, Modified by Daniel Shaykevich
% September 29, 2024 with the help of chatGPT to plot on a logarithmic
% scale. 
locationGroups = cell(1,3);
intervalGroups = cell(1,3);


for i = 1:3
    [fname, pname] = uigetfile('*.*', 'Load the spike location file for next PC');
    cd(pname)
    locationGroups{1,i} = load(fname);    
end

figure
for i = 1:3
    intervalGroups{1,i} = diff(locationGroups{1,i}(:)) * 1000; % in ms

    % Define logarithmic bin edges (for example, from 1 ms to the max interval)
    minBin = 1;  % Starting point for bins (e.g., 1 ms)
    maxBin = max(intervalGroups{1,i});  % Maximum value in the ISI data
    numBins = 50;  % Number of bins (you can adjust this)

    % Generate logarithmic bin edges using logspace
    logBins = logspace(log10(minBin), log10(maxBin), numBins);

    % Create a histogram using the logarithmic bins
    [counts, edges] = histcounts(intervalGroups{1,i}, logBins);
    binCenters = edges(1:end-1) + diff(edges) / 2; % Compute bin centers

    % Plot the bars with equal bin widths on a logarithmic x-axis
    figure
    bar(binCenters, counts, 'histc');  % 'histc' to plot with equal bin width

    set(gca, 'XScale', 'log')  % Set x-axis to log scale (base 10)
    xlabel('Time Interval [ms]', 'fontsize', 18, 'FontName', 'Arial', 'FontWeight', 'bold')
    ylabel('Count', 'fontsize', 18, 'FontName', 'Arial', 'FontWeight', 'bold')
    set(gca, 'FontSize', 14)

 
end
```
