## Supplementary material for "Chronic recording of brain activity in awake toads": Matlab scripts: spike_sorter_core_submission.m

```
% Written by Guosong on Feb 20, 2015.
% Modified by Guosong on July 20, 2015.
% Turned into a 2-arg function on July 23, 2015.
% All rights reserved.
% Constraints:
% 1. Threshold: defined as a set value or median/0.6745*4 or a fixed value
% 2. Artifact: defined as a set value (should not exceed)
% 3. Clear Region: duration and amplitude are both set
% 4. Post-spike amplitude: has to decay to a certain threshold post spike
% 5. Number of ripples: minimize post-spike oscillation

function returnPackage=spike_sorter_Guosong_core(fname,pname,startTime,endTime);

%%%%%%%%%%%%%%%%%%%%%%%%%PARAMETER INITIALIZATIONS%%%%%%%%%%%%%%%%%%%%%%%%%
% Change the start and end times for spike sorting
% Values input here are in seconds
startSortingTime=5e-5+startTime; % Note: startSortingTime has to be non-zero. So if one wants to start from 0, put 5e-5 instead.
endSortingTime= endTime;

% Change what channels to inspect
channelOption='selected'; % Available inputs: 'all' or 'selected'
channels= [11]; % Channel indices. Only used in 'selected' mode.

% Time and amplitude constraints places on spike sorting
preSpike=20; % pre-spike time duration is 1 ms (20/20000)
postSpike=40; % post-spike time duration is 2 ms (40/20000)
clearRegion=1; % 1.0 ms interval required between two neighboring spikes
detectMode='negative'; % Detect positive only, negative only or both signs of peaks
thresholdMode='SetValue'; % Available inputs: 'SetValue' or 'Median' 
mechNoiseSuppress='no'; % For single-batch data analysis without mech noise
thresholdSetValue = 20; % in uV. Only used in 'SetValue' mode.
artifactCutoff = 100; % Maximum amplitude for artifact removal, in uV.

% Bandpass filter initialization
sampleRate=20000; % Hz
passBand=[250 6000]; % Hz
filterOrder=1; 
filterIter=11;
ftype='bandpass';
filterName='butterworth'; % butterworth, elliptic or chebyshev
%%%%%%%%%%%%%%%%%%%%%%%%%%%%%%%%%%%%%%%%%%%%%%%%%%%%%%%%%%%%%%%%%%%%%%%%%%%

cd(pname);
fnamePrefix=fname(1:size(fname,2)-4);
originalData=read_Intan_512RHD_no_prompt(fname,pname); % rows: channels; columns: times
%originalData=read_Intan_RHD2000_file_no_prompt(fname,pname); % rows: channels; columns: times
%%%%%%%%%%%%%%%%%%%%%%%%%%%%% CHANNEL REORDERING %%%%%%%%%%%%%%%%%%%%%%%%%%
% dataBuffer=originalData;
% originalData=[];
% originalData(1,:)=dataBuffer(2,:);
% originalData(2,:)=dataBuffer(1,:);
% originalData(3,:)=dataBuffer(3,:);
% originalData(4,:)=dataBuffer(4,:);
% originalData(5,:)=dataBuffer(6,:);
% originalData(6,:)=dataBuffer(5,:);
% originalData(7,:)=dataBuffer(7,:);
% originalData(8,:)=dataBuffer(8,:);
% originalData(9,:)=dataBuffer(9,:);
% originalData(10,:)=dataBuffer(10,:);
% originalData(11,:)=dataBuffer(25,:);
% originalData(12,:)=dataBuffer(12,:);
% originalData(13,:)=dataBuffer(13,:);
% originalData(14,:)=dataBuffer(14,:);
% originalData(15,:)=dataBuffer(15,:);
% originalData(16,:)=dataBuffer(16,:);
% originalData(1,:)=dataBuffer(1,:);
% originalData(2,:)=dataBuffer(2,:);
% originalData(3,:)=dataBuffer(17,:);
% originalData(4,:)=dataBuffer(18,:);
% originalData(5,:)=dataBuffer(19,:);
% originalData(6,:)=dataBuffer(20,:);
% originalData(7,:)=dataBuffer(21,:);
% originalData(8,:)=dataBuffer(22,:);
% originalData(9,:)=dataBuffer(23,:);
% originalData(10,:)=dataBuffer(24,:);
% originalData(11,:)=dataBuffer(26,:);
% originalData(12,:)=dataBuffer(27,:);
% originalData(13,:)=dataBuffer(28,:);
% originalData(14,:)=dataBuffer(31,:);
% originalData(15,:)=dataBuffer(32,:);
% originalData(16,:)=dataBuffer(8,:);
%%%%%%%%%%%%%%%%%%%%%%%%%%%%%%%%%%%%%%%%%%%%%%%%%%%%%%%%%%%%%%%%%%%%%%%%%%%

if startTime==endTime
    startSortingTime=1/sampleRate;
    endSortingTime=size(originalData,2)/sampleRate;
end
switch channelOption
    case 'selected'
        transposedData=originalData(channels,round(startSortingTime*sampleRate):round(endSortingTime*sampleRate))'; % rows: times; columns: channels
    case 'all'
        transposedData=originalData(:,round(startSortingTime*sampleRate):round(endSortingTime*sampleRate))'; % rows: times; columns: channels
    % Note: 'round' is used here to avoid precision carry error.
end
[numPoints numChannels]=size(transposedData);
t=(startSortingTime:1/sampleRate:endSortingTime)';

% Filter the original traces and save both unfiltered and filtered traces
filteredData=transposedData;
Wn=2*passBand./sampleRate;  % Normalized cutoff frequency. Unitless. 
[butterB,butterA]=butter(filterOrder,Wn,ftype);    %prepare the butterworth filter.
[ellipB,ellipA]=ellip(filterOrder,0.1,100,Wn,ftype);  %prepare the elliptical filter (2nd par: ripple amp; 3rd par: stopband atten).
[chebyB,chebyA]=cheby2(filterOrder,18,Wn,ftype);  %prepare the type-2 Chebyshev filter. (2nd par: stopband atten).
for i=1:filterIter
    switch filterName
        case 'butterworth'
            filteredData=filtfilt(butterB,butterA,filteredData);
        case 'elliptic'
            filteredData=filtfilt(ellipB,ellipA,filteredData);
        case 'chebyshev'
            filteredData=filtfilt(chebyB,chebyA,filteredData);
    end        
end

switch mechNoiseSuppress
    case 'yes'
        for chIndex=1:size(filteredData,2)
            seedIndices=find(abs(filteredData(:,chIndex))>=artifactCutoff);
            grownIndices=seedIndices;
            for offsetIndex=-20:1:20
                offsetIndices=grownIndices+offsetIndex;
                offsetIndices=offsetIndices(find(offsetIndices>0));
                offsetIndices=offsetIndices(find(offsetIndices<=size(filteredData,1)));
                grownIndices=union(grownIndices,offsetIndices);
            end
            filteredData(grownIndices,chIndex)=0;
        end
end

returnPackage=struct('fnamePrefix',fnamePrefix);
returnPackage.unfilteredData=transposedData;
returnPackage.filteredData=filteredData;
returnPackage.time=t;
returnPackage.sampleRate=sampleRate;
returnPackage.numChannels=numChannels;
returnPackage.noise=zeros(numChannels,1);

% Sort out spikes
returnPackage.spikesGroup=cell(numChannels,1);
returnPackage.locationsGroup=cell(numChannels,1);
for traceIndex=1:numChannels
    % Prepare the data for each channel
    filteredChannel=filteredData(:,traceIndex)'; % This once again becomes a single row vector.
    numSelectedSortingRegion=size(filteredChannel,2);
    
    % Set threshold for detection of spikes
    switch thresholdMode
        case 'SetValue'
            threshold=thresholdSetValue;
        case 'Median'
            threshold=median(abs(filteredChannel))/0.6745*4;
    end
    returnPackage.noise(traceIndex)=threshold/4;
    
    % Locate the spike times
    switch detectMode
        case 'positive'
            index = find(filteredChannel(preSpike+clearRegion+1:end-postSpike-clearRegion) > threshold) +preSpike+clearRegion;
        case 'negative'
            index = find(filteredChannel(preSpike+clearRegion+1:end-postSpike-clearRegion) < -threshold) +preSpike+clearRegion;
        case 'both'
            index = find(abs(filteredChannel(preSpike+clearRegion+1:end-postSpike-clearRegion)) > threshold) +preSpike+clearRegion;
    end

    % Converge data points for one spike
    index_length=length(index);
    total_data_pts=preSpike+postSpike+1;
    spikes=zeros(index_length,total_data_pts);
    for i=1:index_length                          
        maxAbsValue=max(abs(filteredChannel(index(i)-preSpike-clearRegion:index(i)+postSpike+clearRegion)));
        if  maxAbsValue<artifactCutoff 
            switch detectMode
                case 'positive'
                    maxValue=0;
                    while max(filteredChannel(index(i)-preSpike:index(i)+postSpike))>maxValue && max(abs(filteredChannel(index(i)-preSpike:index(i)+postSpike)))<artifactCutoff && (index(i)-2*preSpike-clearRegion)>0 && (index(i)+2*postSpike+clearRegion)<=numSelectedSortingRegion
                        maxValue=max(filteredChannel(index(i)-preSpike:index(i)+postSpike));
                        index(i)=find(filteredChannel(index(i)-preSpike:index(i)+postSpike)==maxValue)+index(i)-preSpike-1;
                    end
                case 'negative'
                    minValue=0;
                    while min(filteredChannel(index(i)-preSpike:index(i)+postSpike))<minValue && max(abs(filteredChannel(index(i)-preSpike:index(i)+postSpike)))<artifactCutoff && (index(i)-2*preSpike-clearRegion)>0 && (index(i)+2*postSpike+clearRegion)<=numSelectedSortingRegion
                        minValue=min(filteredChannel(index(i)-preSpike:index(i)+postSpike));
                        index(i)=find(filteredChannel(index(i)-preSpike:index(i)+postSpike)==minValue)+index(i)-preSpike-1;
                    end
                case 'both'
                    maxValue=0;
                    while max(abs(filteredChannel(index(i)-preSpike:index(i)+postSpike)))>maxValue && max(abs(filteredChannel(index(i)-preSpike:index(i)+postSpike)))<artifactCutoff && (index(i)-2*preSpike-clearRegion)>0 && (index(i)+2*postSpike+clearRegion)<=numSelectedSortingRegion
                        maxValue=max(abs(filteredChannel(index(i)-preSpike:index(i)+postSpike)));
                        index(i)=find(abs(filteredChannel(index(i)-preSpike:index(i)+postSpike))==maxValue)+index(i)-preSpike-1;
                    end
            end
            maxClearRegion=max(max(abs(filteredChannel(index(i)-preSpike-clearRegion:index(i)-preSpike-1))),max(abs(filteredChannel(index(i)+postSpike+1:index(i)+postSpike+clearRegion))));
            mainPeak=filteredChannel(index(i));
            maxPostSpike=max(abs(filteredChannel(index(i)+postSpike/2:index(i)+postSpike)));
            maxPreSpike=max(abs(filteredChannel(index(i)-preSpike:index(i)-preSpike*0.4)));
            if mainPeak>0
                if maxClearRegion<mainPeak*0.5 && mainPeak<artifactCutoff && maxPostSpike<mainPeak*0.4 && maxPreSpike<mainPeak*0.8
                    spikes(i,:)=filteredChannel(index(i)-preSpike:index(i)+postSpike);
                end
            else
                if maxClearRegion<abs(mainPeak)*0.5 && abs(mainPeak)<artifactCutoff && maxPostSpike<abs(mainPeak)*0.4 && maxPreSpike<abs(mainPeak)*0.8
                    spikes(i,:)=filteredChannel(index(i)-preSpike:index(i)+postSpike);
                end
            end
        end
    end
   % artifacts = find(spikes(:,preSpike)==0);       % Erase indices that were artifacts
   % spikes(artifacts,:)=[];
   % index(artifacts)=[];

    % Clean up redundancy
    [index sortNumber]=sort(index);
    spikes=spikes(sortNumber,:);
    currentIndex=0;
    toRemoveIndex=[];
    for i=1:size(index,2)
        if abs(index(i)-currentIndex)<postSpike+clearRegion
            toRemoveIndex=[toRemoveIndex i];
        else
            currentIndex=index(i);
        end
    end
    spikes(toRemoveIndex,:)=[];
    index(toRemoveIndex)=[];
    
    [numSpikes numDataPts]=size(spikes);
    
    returnPackage.spikeTimeStamp=([1:numDataPts]*1000/sampleRate)';
    returnPackage.spikesGroup{traceIndex}=spikes';
    returnPackage.indicesGroup{traceIndex}=index';
end
```
