## Supplementary material for "Chronic recording of brain activity in awake toads": Matlab scripts: spike_sorter_submission.m

```
% Written by Guosong on Feb 20, 2015.
% Modified by Guosong on July 20, 2015.
% Modified by Daniel Shaykevich on October 6th, 2024. Adjusted this script with 
% the help of chatGPT, to plot mean waveforms with standard deviation shading, 
% and checked the work to confirm it was accurate
% All rights reserved.
% Constraints:
% 1. Threshold: defined as a set value or SD*3 or SD*5
% 2. Artifact: defined as a set value (should not exceed)
% 3. Clear Region: duration and amplitude are both set
% 4. Post-spike amplitude: has to decay to a certain threshold post spike
% 5. Number of ripples: minimize post-spike oscillation

% Initialization
clear;
close all;
set(0,'DefaultFigureWindowStyle','docked')

%%%%%%%%%%%%%%%%%%%%%%%%%%PARAMETER INITIALIZATION%%%%%%%%%%%%%%%%%%%%%%%%%
% Multi-channel plot offsets
unfilteredPlotOffset=2000;
filteredPlotOffset= 15;
%%%%%%%%%%%%%%%%%%%%%%%%%%%%%%%%%%%%%%%%%%%%%%%%%%%%%%%%%%%%%%%%%%%%%%%%%%%

%%%%%%%%%%%%%%%%%%%%%%%%%%%%%%%%FILE I/O%%%%%%%%%%%%%%%%%%%%%%%%%%%%%%%%%%%
% File input
[fname pname]=uigetfile('*.*','Please select the Intan file to open');

% File output
pnameToSave=uigetdir(pname,'Please select the folder to save the processed files');
%%%%%%%%%%%%%%%%%%%%%%%%%%%%%%%%%%%%%%%%%%%%%%%%%%%%%%%%%%%%%%%%%%%%%%%%%%%

results=spike_sorter_core_submission(fname, pname,0, 60);

%%%%%%%%%%%%%%%%%%%%%%%%%%%%%LOADING FIELDS%%%%%%%%%%%%%%%%%%%%%%%%%%%%%%%%
fnamePrefix=results.fnamePrefix;
unfilteredData=results.unfilteredData;
filteredData=results.filteredData;
time=results.time;
sampleRate=results.sampleRate;
numChannels=results.numChannels;
spikeTimeStamp=results.spikeTimeStamp;
spikesGroup=results.spikesGroup;
indicesGroup=results.indicesGroup;
noise=results.noise;
%%%%%%%%%%%%%%%%%%%%%%%%%%%%%%%%%%%%%%%%%%%%%%%%%%%%%%%%%%%%%%%%%%%%%%%%%%%

cd(pnameToSave);
unfilteredFileName=strcat(fnamePrefix,'_unfiltered.dat');
filteredFileName=strcat(fnamePrefix,'_filtered.dat');
noiseFileName=strcat(fnamePrefix,'_noise.dat');
command=sprintf('save %s unfilteredData -ascii', unfilteredFileName);
eval(command);
command=sprintf('save %s filteredData -ascii', filteredFileName);
eval(command);
command=sprintf('save %s noise -ascii', noiseFileName);
eval(command);

%Plot individual traces
for traceIndex=1:numChannels
    figure
    hold on
    for i=1:1:size(spikesGroup{traceIndex},2)
        if filteredData(indicesGroup{traceIndex}(i),traceIndex)>0
            plot(spikeTimeStamp,spikesGroup{traceIndex}(:,i),'r-','Linewidth',2);
        else plot(spikeTimeStamp,spikesGroup{traceIndex}(:,i),'b-','Linewidth',2);
        end
        hold on
    end
    xlabel('Time [ms]','fontsize',36,'FontName','Arial','FontWeight','bold')
    ylabel('Voltage [uV]','fontsize',36,'FontName','Arial','FontWeight','bold')
    set(gca,'FontSize',24,'Linewidth',2,'box','off')
    axis([0 3 -50 50])
    toSaveMat=[spikeTimeStamp spikesGroup{traceIndex}];
    indexStr=num2str(traceIndex);
    while size(indexStr,2)<2
        indexStr=strcat('0',indexStr);
    end
    toSaveName=strcat('Spikes_Channel_',indexStr,'.dat');
    command=sprintf('save %s toSaveMat -ascii', toSaveName);
    eval(command);
    
    peakLocationName=strcat('Spike_Locations_Channel_',indexStr,'.dat');
    peakLocation=indicesGroup{traceIndex}/sampleRate+time(1)-1/sampleRate;
    peakIndicator=[peakLocation filteredData(indicesGroup{traceIndex},traceIndex)+((filteredData(indicesGroup{traceIndex},traceIndex)>0)-0.5)*100];
    command=sprintf('save %s peakLocation -ascii', peakLocationName);
    eval(command);
    figure 
    plot(time,filteredData(:,traceIndex),'k-','Linewidth',2);
    hold on
    plot(peakIndicator(find(peakIndicator(:,2)>0),1),peakIndicator(find(peakIndicator(:,2)>0),2),'r*');
    plot(peakIndicator(find(peakIndicator(:,2)<0),1),peakIndicator(find(peakIndicator(:,2)<0),2),'b*');
   % axis([time(1) time(size(time,1)) -200 200]) 
   axis([0 60 -200 200]) 
    xlabel('Time [s]','fontsize',36,'FontName','Arial','FontWeight','bold')
    ylabel('Voltage [uV]','fontsize',36,'FontName','Arial','FontWeight','bold')
    set(gca,'FontSize',24,'Linewidth',2,'box','off')
end

% Plot mean waveform with sd shading (commented out range shading)
for traceIndex = 1:numChannels
    % Extract spike waveforms for the current channel
    spikes = spikesGroup{traceIndex};
    
    % Find non-zero columns
    nonZeroCols = any(spikes, 1);  % Identify columns with non-zero values
    spikesNonZero = spikes(:, nonZeroCols);  % Select only non-zero columns
    
    % Time axis in milliseconds (assuming spikeTimeStamp is in ms)
    x = spikeTimeStamp;  % This should match the size of your spikes data

    % Calculate the mean waveform
    meanWaveform = mean(spikesNonZero, 2);  % Mean across rows for non-zero columns
    minWaveform = min(spikesNonZero, [], 2);  % Minimum across rows
    maxWaveform = max(spikesNonZero, [], 2);  % Maximum across rows
    stdWaveform = std(spikesNonZero, 0, 2);
   

    figure;  % Create a new figure
    hold on;

    % Plot the mean waveform with sd
    plot(x, meanWaveform, 'k', 'LineWidth', 2);  % Black line for the mean waveform

    % Fill the area between min and max for each time point (range shading)
    %fill([x; flipud(x)], [maxWaveform; flipud(minWaveform)], 'k', 'FaceAlpha', 0.3, 'EdgeColor', 'none');

    % Fill the area within one standard deviation of the mean
    fill([x; flipud(x)], [(meanWaveform + stdWaveform); flipud((meanWaveform-stdWaveform))], 'k', 'FaceAlpha', 0.3, 'EdgeColor', 'none');

    % Set axis limits based on the spike time and voltage range
    xlim([min(x) max(x)]);  % X-axis is time in ms
    ylim([-200 200]);        % Adjust based on your data's expected voltage range

    % Axis properties
    xlabel('Time [ms]', 'fontsize', 12, 'FontName', 'Arial', 'FontWeight', 'bold');
    ylabel('Voltage [uV]', 'fontsize', 12, 'FontName', 'Arial', 'FontWeight', 'bold');
    title(['Mean Waveform + Voltage Range - Channel ', num2str(traceIndex)], 'fontsize', 12);
    set(gca, 'FontSize', 10, 'LineWidth', 1.5, 'box', 'off');
    
    % Plot appearance
    set(gca, 'XColor', 'k', 'YColor', 'k');
    set(gca, 'TickDir', 'out');
    set(gca, 'TickLength', [0.02, 0.02]);
    box off;
    
    % Optional: Adjust the layout
    sgtitle(['Spike Analysis for Channel ', num2str(traceIndex)], 'fontsize', 14, 'FontWeight', 'bold');
end

% Plot all original, unfiltered traces
figure
for i=1:numChannels
    plot(time,unfilteredData(:,i)+unfilteredPlotOffset*(i-1),'k-');
    hold on
    %axis([time(1) time(size(time,1)) -3*unfilteredPlotOffset unfilteredPlotOffset*(numChannels-1)+unfilteredPlotOffset*3])
    axis([0 60 -3*unfilteredPlotOffset unfilteredPlotOffset*(numChannels-1)+unfilteredPlotOffset*3])
end
set(gca,'FontSize',14,'Linewidth',2)
xlabel('Time [s]','fontsize',18,'FontName','Arial','FontWeight','bold')
ylabel('Voltage [uV]','fontsize',18,'FontName','Arial','FontWeight','bold')
title('Original Data')

figure
for i=1:numChannels
    hold on
    plot(time,filteredData(:,i)+filteredPlotOffset*(i-1),'k-');
    %axis([time(1) time(size(time,1)) -3*filteredPlotOffset filteredPlotOffset*(numChannels-1)+filteredPlotOffset*3])
    axis([0 60 -3*filteredPlotOffset filteredPlotOffset*(numChannels-1)+filteredPlotOffset*3])
end
set(gca,'FontSize',14,'Linewidth',2)
xlabel('Time [s]','fontsize',18,'FontName','Arial','FontWeight','bold')
ylabel('Voltage [uV]','fontsize',18,'FontName','Arial','FontWeight','bold')
title('Filtered Data')
```
